# Allo-Allo: Data-efficient prediction of allosteric sites

**DOI:** 10.1101/2024.09.28.615583

**Authors:** Tianze Dong, Christopher Kan, Kapil Devkota, Rohit Singh

**Affiliations:** Department of Computer Science, Duke University; Department of Mathematics, Duke University; Department of Biostat. and Bioinformatics, Duke University; Department of Biostat. and Bioinformatics, Department of Cell Biology, Duke University

## Abstract

Allostery, a fundamental structural mechanism where ligand binding at a protein site affects protein function at another site, plays a crucial role in key drug-target proteins like GPCRs. Unfortunately, existing methods for predicting allosteric sites have limited performance– they are particularly constrained by scarce ground-truth experimental data. We introduce Allo-Allo, a data-efficient, sequence-based method that predicts allosteric sites by leveraging protein language models (PLMs). Honing in on ESM-2 attention heads that capture allosteric residue associations, Allo-Allo achieves a 67% higher AUPRC than state-of-the-art methods. Our innovative, data-efficient pipeline not only outperforms alternate, commonly-used PLM-based prediction architectures but also generalizes well. Notably, mutations in Allo-Allo-predicted sites show significant association with elevated disease risk scores from AlphaMissense, highlighting its translational potential. Beyond Allo-Allo’s biological and translational applicability, its architecture presents a powerful framework for other data-scarce problems in protein analysis.

## 1 Introduction

In the post-AlphaFold era, with accurate but static structures now inferable, the field’s limited understanding of protein conformational flexibility and dynamics has emerged as a crucial research gap. A biologically vital form of conformational flexibility is *allostery*: a mechanism by which ligand binding at a particular site (the *allosteric site*) induces conformational changes that impact binding at other sites (e.g., the active or *orthosteric site*). Conceptually, allostery acts as a signal integrator, enabling proteins to respond dynamically to cellular signals and environmental cues. It thus plays a critical regulatory role in diverse biological contexts such as signal transduction, metabolism, and transcriptional regulation [7, 17, 18]. Due to their central role in many biological processes, allosteric proteins are prime targets for drug interventions. For instance, G-protein coupled receptors (GPCRs) have been popular drug targets, with over 30% of FDA-approved small-molecule drugs targeting them [10]. Allostery is key to their functioning, and further therapeutic progress with GPCRs hinges on accurately identifying and targeting their allosteric sites.

Here we focus on the critical first step in understanding allostery: predicting the allosteric site of a protein. Unfortunately, current approaches to predicting allosteric sites either do not scale beyond a subset of proteins or have limited accuracy. Techniques such as PARS [14] and SPACER [9] apply molecular dynamics (MD) to locate allosteric binding pockets that induce conformational changes in the protein. However, the applicability of MD-based methods is limited by the scarcity of reliable, long-duration MD data. An alternative set of approaches is information theoretic, reasoning that some latent structural mechanism should link the allosteric and orthosteric sites. The state-of-the-art PASSer family of models [22, 24, 26], operates on a single, static structural conformation and uses a variety of methods including XGBoost and message-passing graph neural networks to identify allosteric sites. However, the performance of such data-driven approaches has been limited.

The fundamental challenge faced by existing methods like PASSer is the paucity of experimental data on allostery. Ground truth data on allosteric sites is typically obtained by low-throughput mutational studies and is currently available only for a limited set of proteins. The problem is compounded by poor data annotation. The AlloSteric Database (“ASD”; version 5.1, 2023) [11] reports around 2,422 proteins with known allosteric sites. Our analysis of the database entries revealed inconsistencies between the sequence information from reference sources such as PDB [3] and UniProt [21] and the position-specific residues listed in ASD. After discarding non-matching entities, only 653 well-annotated proteins were usable for computational purposes. This data limitation underscores the need for a model that can effectively train on limited data while achieving strong predictive performance.

To design an accurate data-efficient method for predicting allosteric sites, we leverage protein language models (PLMs) [2, 16, 8, 19, 12]. Since allostery is such an important biological phenomenon that it is evolutionarily conserved [27, 4], we hypothesized that foundation-model PLMs, trained on millions of protein sequences, must already be capturing it. The key conceptual advance of our work is that rather than using the limited ground-truth data on allostery to train a predictive model from scratch, we instead use it to identify the parts of a pretrained PLM’s internal representation that already capture allostery and train a simple but performant model on top of that. We introduce Allo-Allo for predicting allosteric sites solely from sequence. We reason that latent structural connections between allosteric site and other parts of the protein can be discerned through internal PLM representations. Specifically, we identify the attention heads in ESM-2 PLM models that capture the association between allosteric sites and the rest of the protein, building upon the model-interrogation framework of Vig et al. [25]. The selected attention scores are then combined in a random forest (RF) predictor.

Allo-Allo strongly outperforms existing approaches, achieving an AUPRC of 0.77 on allosteric site prediction, compared to the current best AUPRC of 0.46 (PASSerRank). This outperformance is especially notable considering that Allo-Allo only requires sequence information while the PASSer family of methods require protein structure as input. Remarkably, Allo-Allo achieves strong allosteric site prediction even without knowledge of orthosteric sites, as we show in an ablation study. Our findings suggest that allosteric sites play an overall critical role in conformational dynamics, not just limited to orthosteric sites. To explore this further, we cross-referenced our allosteric site predictions on 2,567 cell-surface proteins against variant-risk scores from AlphaMissense [5], finding substantial concordance. We expect Allo-Allo’s conceptual innovations for data-efficient prediction will be broadly applicable and its predictions of allosteric sites will be valuable during drug discovery.

## 2 Methods

### Data curation and pre-processing

We curated data from the AlloSteric Database (ASD), which contained 2,422 proteins with annotated allosteric sites. After cross-referencing with the PDB and UniProt to ensure consistency, there remained only 653 proteins with well-matched annotations. We split this dataset into training, validation, and test sets in a 70:10:20 ratio. The training and validation sets were used for ablation studies to fine-tune model parameters. After completing the ablations, we combined these two sets to fit a final model, which was then benchmarked against other methods and applied for disease risk prediction. In the RF classifier, residues not labeled as allosteric sites within these proteins were treated as negative examples.

### Quantifying allosteric sensitivity in attention heads

Assume that the PLM contain *L* layers, each with *H* heads (e.g., *l* = 32 and *H* = 20 for the 650M-parameter ESM-2 model). Given a protein sequence ***x*** = (*x*_1_, …, *x*_*n*_) of length *n*, each attention head, ***A***_***ℓ***,***h***_ *∈* ℝ^*n×n*^, *ℓ ∈* 1, …, *L, h ∈* 1, …, *H*, is a *n × n* matrix encoding pairwise residue relationships.

We followed Vig et al.’s [25] approach to score attention heads. Suppose *S*_*allo*_ = {*α*_1_, …, *α*_*k*_} represent the set of known *k* allosteric sites of ***x***. Then, for all attention heads, we calculate how strong the pairwise associations are between the locations in *S*_*allo*_ and the rest of the sequence. Setting *θ* as the minimum score needed for the pairwise residue relationships to be considered significant, the allosteric activity for (*ℓ, h*), given by *w*_*allo*_(***A***_***ℓ***,***h***_(***x***)), was computed the following way:

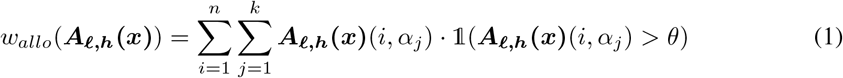

The total activity of the attention head ***A***_***ℓ***,***h***_, *w*(***A***_***ℓ***,***h***_**(*x*)**), is then computed as:

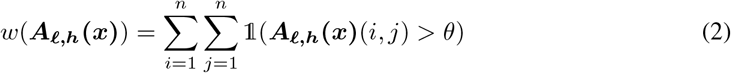

The allosteric impact, i.e. the fraction of total activity in ***A***_***ℓ***,***h***_ contributed by allostery, is 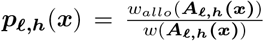For all training sequences, *x*^(1)^, …, *x*^(*N*)^, the resulting set 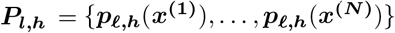 was constructed for all *ℓ* and *h* pairs.

### Selecting attention heads with highest allosteric sensitivity

In the second step, we applied the Student’s t-test to each ***P***_***ℓ***,***h***_ set, with the null hypothesis implying a *µ*_**0**_ equal to the average *p*-score across all training samples and attention heads. Bonferroni correction was applied to adjust the *p*-values for multiple hypothesis testing. Finally, the attention heads that rejected the null hypothesis were ranked based on their Signal-to-Noise Ratios (SNRs), where 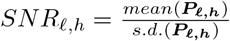 We chose *K* attention heads with the largest SNR values for subsequent feature selection.

### Using selected attention heads to train the Random Forest model

After finding the *K* allostery-sensitive attention-heads, we next generated features for each residue *x*_*i*_ in an input sequence ***x*** by row-averaging the selected *K* attention heads at their *i*^*th*^ rows. For each residue, this resulted in a vector of *K* features which, along with the ground truth residue allostery label, was used to train a RF classifier. All the residues in the training set were used during training; the residues that were not marked as allosteric were labelled as negative training examples.

### Alternative approaches

We considered two alternative approaches and performed ablation studies for them. In the first approach, we simply trained a multi-layer perceptron (MLP) classifier on top of fixed ESM-2 650M embeddings. The second approach is essentially the same as Allo-Allo, except that attention weights were quantified only on orthosteric-allosteric pair interactions (**Appendix B**).

#### Related work

Previously, data driven approaches like PASSer have been shown to outperform MD-based approaches [22]. The state of the art comprises the **PASSer** family of models [26, 22, 24, 23]. They seek to identify the latent structural links between allosteric and orthosteric sites from protein structure, offering three distinct architectures: (a) XGBoost-based PASSer, (b) AutoGluon-based PASSer2.0 and (c) PASSerRank, that applies the LambdaMART [13] algorithm to rank potential allosteric sites. Additionally, we also compare to the SVM-based AllositePro [20] algorithm.

## 3 Results

### Benchmarking

Across a wide range of metrics, Allo-Allo significantly outperformed existing methods (**Table 1**). Interestingly, Allo-Allo’s attention head strategy substantially outperformed an alternate prediction head-based approach, widely used in PLM literature (blue row in **Table 1**; **Appendix A**). The outperformance of Allo-Allo underscores that the intrinsic residue relationships contained within attention heads were already learned by foundation models. Our novel pipeline, employing t-tests and SNR scoring along with RF-based classification, effectively prioritizes and leverages these attention heads to extract more signal from the limited ground-truth data than the commonly-used prediction head-based design.

**Table 1:**
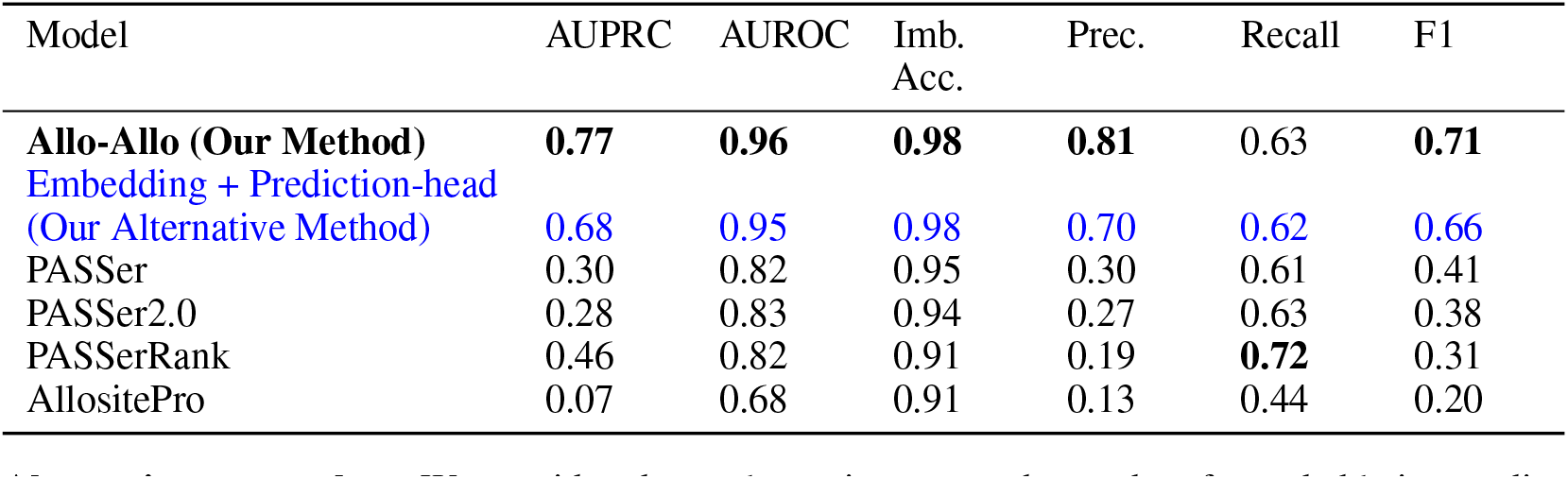
Comparison of our method with state-of-the-art models on allosteric sites prediction.

| Model | AUPRC | AUROC | Imb.<br>Acc. | Prec. | Recall | F1 |
| --- | --- | --- | --- | --- | --- | --- |
| **Allo-Allo (Our Method)** | **0.77** | **0.96** | **0.98** | **0.81** | 0.63 | **0.71** |
| **Embedding + Prediction-head**<br>**(Our Alternative Method)** | **0.68** | **0.95** | **0.98** | **0.70** | **0.62** | **0.66** |
| PASSer | 0.30 | 0.82 | 0.95 | 0.30 | 0.61 | 0.41 |
| PASSer2.0 | 0.28 | 0.83 | 0.94 | 0.27 | 0.63 | 0.38 |
| PASSerRank | 0.46 | 0.82 | 0.91 | 0.19 | **0.72** | 0.31 |
| AllositePro | 0.07 | 0.68 | 0.91 | 0.13 | 0.44 | 0.20 |

Another design choice we explored was whether to prioritize information on orthosteric sites during attention computation. While previous work has required knowledge of orthosteric sites, Allo-Allo does not, giving it a usability advantage. In an ablation, we evaluated an Allo-Allo variant focused primarily on orthosteric-allosteric pairwise interaction during attention-head selection and feature extraction (**Appendix B**). We found that this variant performed substantially worse (**Supplement Table A1**). Our findings are concordant with previous reports [4, 27] that allosteric sites have a global impact on protein conformational dynamics, beyond just the orthosteric sites. Moreover, our results suggest that this evolutionary pressure is captured by ESM-2’s attention heads.

### Other ablation studies

We additionally performed ablation on three key criteria important for Allo-Allo feature selection: a) the size of the ESM-2 PLM, b) the threshold *θ* used for quantifying for allosteric impact, and c) number of attention heads (*K*) selected for featurization (**Table 2**).

**Table 2:**
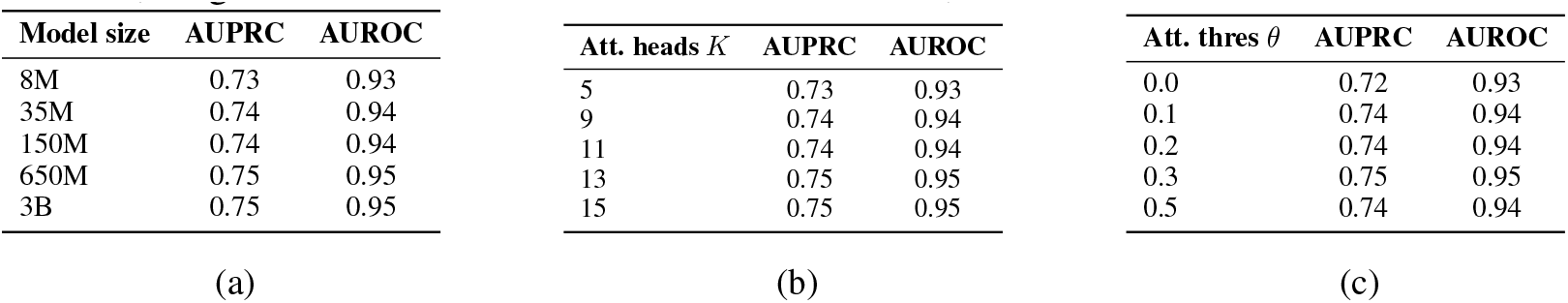
Ablation studies: (a) Impact of ESM-2 Model Size on Performance Metrics (with 15 attention heads and *θ* = 0.3), (b) Impact of the Number of Attention Heads on Performance Metrics (using ESM-2 650M model and *θ* = 0.3), and (c) Impact of Attention Threshold *θ* on Performance Metrics (using ESM-2 650M model and 15 attention heads).

| Model size | AUPRC | AUROC | Att. heads $K$ | AUPRC | AUROC | Att. thres $\theta$ | AUPRC | AUROC |
| --- | --- | --- | --- | --- | --- | --- | --- | --- |
| 8M | 0.73 | 0.93 | 5 | 0.73 | 0.93 | 0.0 | 0.72 | 0.93 |
| 35M | 0.74 | 0.94 | 9 | 0.74 | 0.94 | 0.1 | 0.74 | 0.94 |
| 150M | 0.74 | 0.94 | 11 | 0.74 | 0.94 | 0.2 | 0.74 | 0.94 |
| 650M | 0.75 | 0.95 | 13 | 0.75 | 0.95 | 0.3 | 0.75 | 0.95 |
| 3B | 0.75 | 0.95 | 15 | 0.75 | 0.95 | 0.5 | 0.74 | 0.94 |

We evaluated ESM-2 models of varying scale, with parameter counts ranging from 8M to 3B. We found that the allosteric property is largely captured across all model sizes. In fact, the 650M model demonstrated performance comparable to the 3B version, indicating that increasing the model size beyond 650M parameters does not necessarily enhance performance. This aligns with previous findings that, after a point, scaling PLM models does not enhance their expressiveness of a protein’s functional attributes [15]. It is also notable that even the 8M model does well, pointing to allostery’s strong evolutionary conservation.

We also found that the performance remains largely stable across a wide range of the key Allo-Allo parameters, *K* and *θ*. We found that the selected heads were primarily from the later layers, agreeing with previous findings that early PLM layers capture basic relationships while later layers capture more complex structural properties. Interestingly, even selecting just 5 attention heads for featurization results in only a minor reduction of Allo-Allo’s inference abilities. For the final model selection, we trained the Allo-Allo model by setting *K* = 15 and *θ* = 0.3.

### Allo-Allo’s predicted allosteric sites are associated with elevated risk of disease

To explore the potential of Allo-Allo in biological research, we applied it to score allosteric site probability for 2,567 human proteins from the Surfaceome database [1]. The set comprises proteins (including GPCRs) that reside on the cell surface. Not only are such proteins translational targets, they often play a significant role in signal transduction.

We reasoned that mutations in allosteric sites of these proteins could disrupt their function, increasing the risk of disease. Accordingly, we compared Allo-Allo allosteric site predictions with AlphaMissense estimates of per-residue disease risks. The latter were generated from Cheng et al.’s state-of-the-art deep learning model trained on genetic variant data. The AlphaMissense corpus provides pathogenicity risk scores for the 19 potential residue substitutions at each location of all human proteins [6]. We evaluated three strategies to aggregate these into a single per-location score: (i) average or the (ii) maximum of the 19 scores at each location. Additionally, Cheng et al. also specified a pathogenicity threshold, which we used to compute a (iii) weighted risk-score (**Figure 2A**; **Appendix C**).

**Figure 1.**
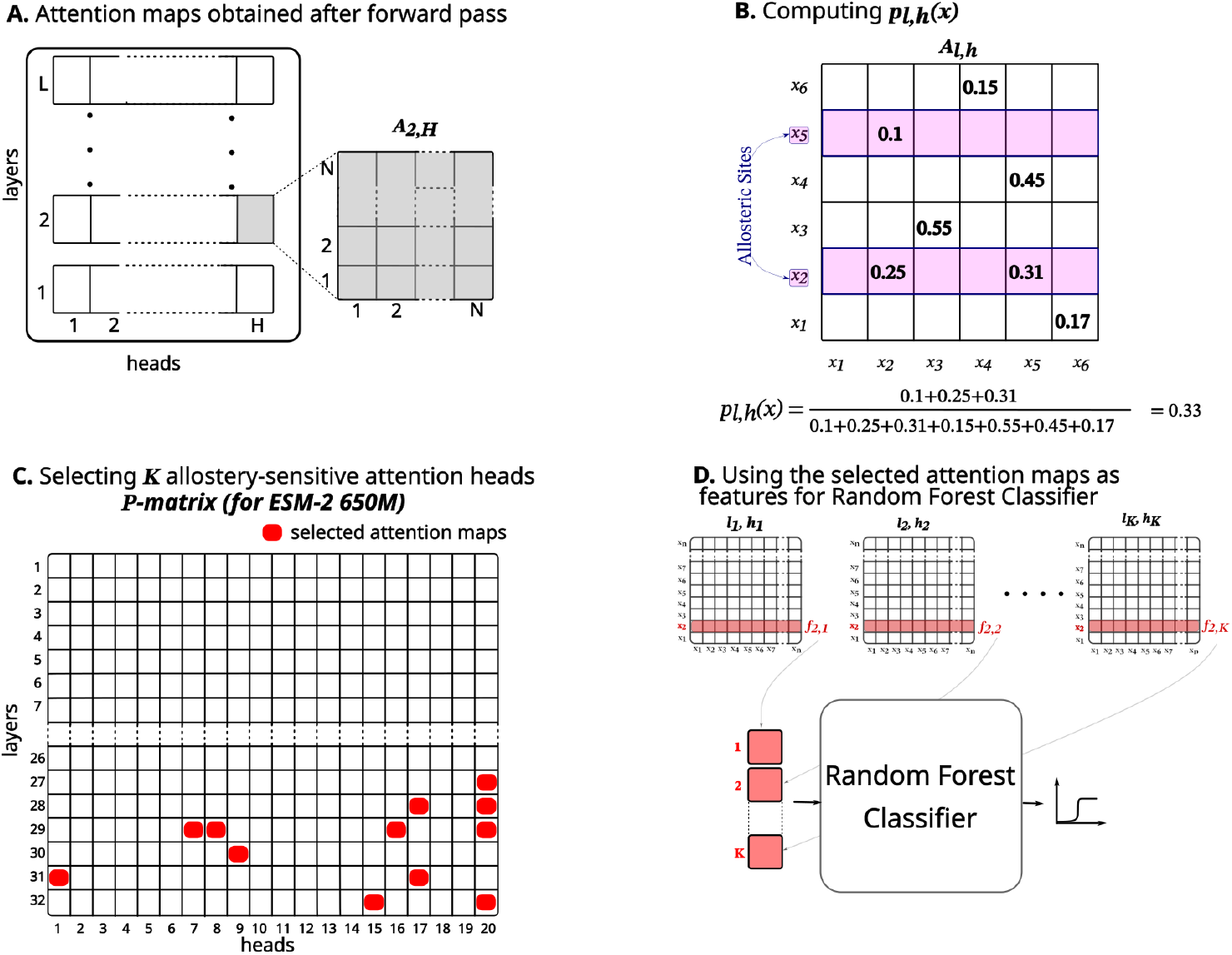
Allo-Allo pipeline: **A**.Organization of the PLM attention heads in a forward operation; each of *L* transformer layers contains *H* attention heads, each head being a *N × N* matrix of pairwise residue relationships. **B**. Allo-Allo computes allosteric impact by computing the degree of contribution from allosteric sites to the overall signal. **C**. The allosteric impact is used to select allostery sensitive attention heads **D**. Their information is used to featurize residues for a RF classifier.

**Figure 2.**
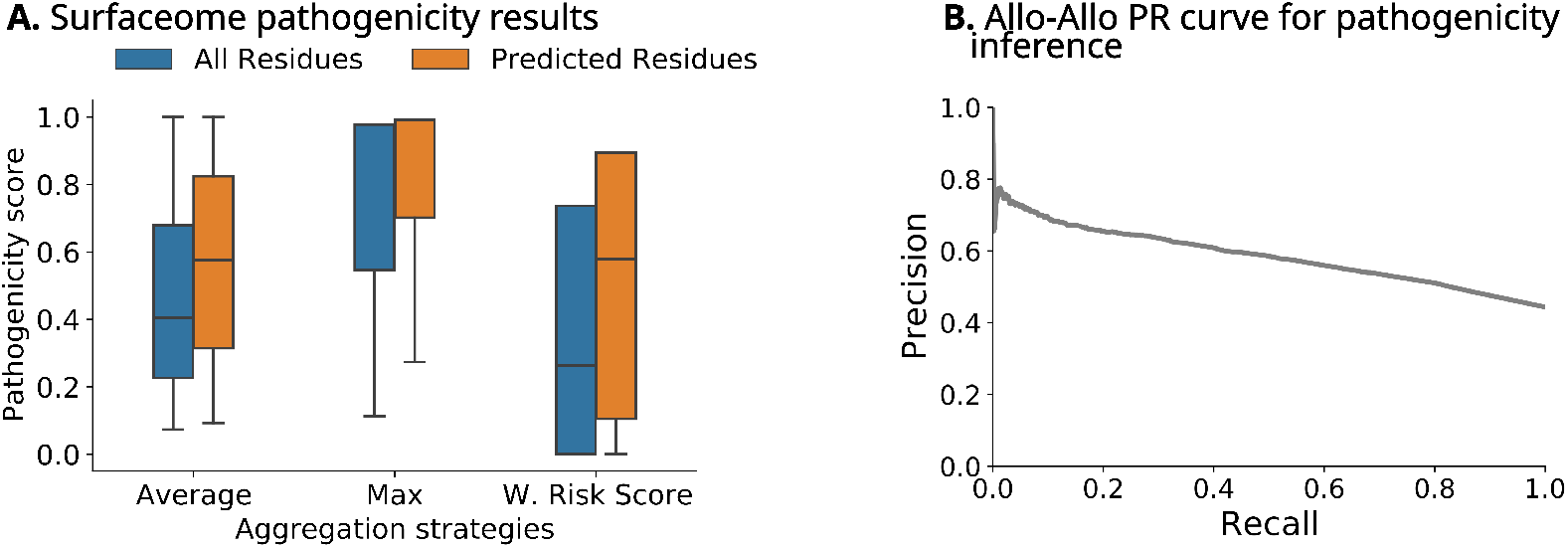
Comparing with AlphaMissense pathogenicity scores: **A**.Pathogenicity aggregated per residue, compared between all analyzed residues and Allo-Allo predicted residues. **B**. Precision-Recall Curve of Allo-Allo probability values as a predictor of a residue’s pathogenicity.

Allo-Allo’s predictions show strong agreement with AlphaMissense risk scores. We marked Allo-Allo predictions above the 99-th percentile as positive, the rest as negative, and compared the distribution of AlphaMissense risk-scores between the two sets. Through a Welch’s t-test, we found that the Allo-Allo predictions had significantly higher scores in the average (*t* = 9.46), max (*t* = 9.01), and weighted risk-score (*t* = 8.86); *p <* 10^*™*5^ in all cases. As an alternative assessment, we labeled the 20% least-risky AlphaMissense sites as negative, the 20% most-risky sites as positive, and evaluated how well the Allo-Allo probabilities separated the two classes. The classifier obtained an AUPRC of 0.59 (**Figure 2B**), substantially better than random (=0.44).

## Discussion

The fundamental importance of allostery is reflected in PLMs, with even the smallest ESM-2 model (8M) capturing it surprisingly well. Just like long-range contacts, allosteric interactions are captured primarily by the later ESM-2 layers, pointing to their complexity. After prioritizing the attention heads, our statistical tests and RF-based pipeline offer a data-efficient approach that may translate to other domains.

Allo-Allo’s concordance with AlphaMissense not only underscores our method’s generalizability but also suggests a valuable application direction in exploring the functional impact of allostery and its role in protein interactions. Future work could also aim to expand the training data available by better curation, or by incorporating MD-based simulations as synthetic data.

## APPENDIX A Appendix

### A Prediction-Head Method

As an alternative to the attention-based Allo-Allo, we explored a baseline method that directly leverages the output ESM-2 representations to predict allosteric sites. This method involves training a simple prediction head, which in our case is a Multi-Layer Perceptron (MLP) classifier, that takes in the ESM-2 embeddings and returns residue-specific allosteric probabilities.

#### Model description

The MLP architecture comprises two linear layers, seperated by a non-linear ReLU. The input dimension *d*_*in*_ = 1280 matches the representation space of ESM-2 650M. The final output returned by the prediction head is a column vector of allosteric probabilities for each residue location. The overall architecture is shown in **Figure A1**.

**Figure A1.**
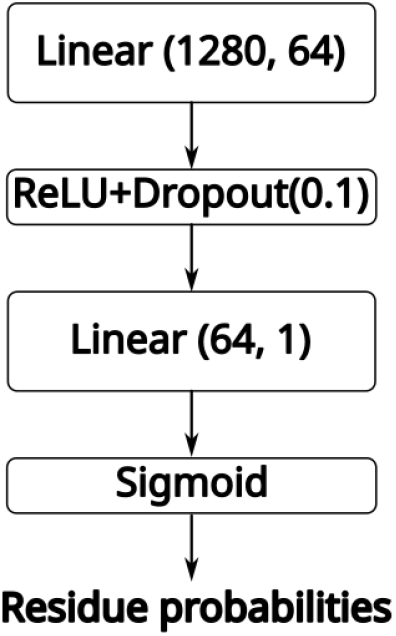
Architecture of the MLP prediction head.

#### Training

The MLP was trained using the binary cross-entropy loss:

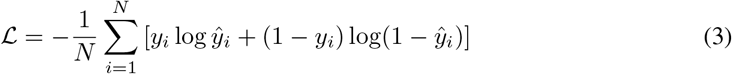

where *N* is the total number of residues in the training set, and *y*_*i*_ *∈* {0, 1} is the ground truth label indicating whether residue *x*_*i*_ is part of an allosteric site.

We used the Adam optimizer with a learning rate of *η* = 1 *×* 10^*™*2^ and implemented early stopping based on validation loss to prevent overfitting. The model was trained for 10 epochs.

#### Results

The Prediction-Head approach achieved an AUPRC of 0.68 and an AUROC of 0.95 on the test set (see Table 1). While higher than the state-of-the-art methods like PASSer, this approach significantly underperformed against attention-head based Allo-Allo. The benchmarking results demonstrate that function-enriched signals contained in attention heads could be harnessed for inference purpose more effectively than the prediction-head strategy, especially in a data-scarce regime.

### B Allo-Allo-Ortho

As an ablation, we evaluated the effects of orthosteric sites in allosteric inference by capitalizing only on the pairwise allosteric-orthosteric relationships contained in the attention heads, ignoring signals coming from non-orthosteric residues. This resulted in a new model, which we named Allo-Allo-Ortho. Unlike the original version, Allo-Allo-Ortho method requires information about the orthosteric sites during the model training and inference.

There were some entries lacking the information about othosteric sites in the ASD. To make our comparisons fairer, we removed those entries from the original set, leaving behind 576 examples. This data was further split into train (80%) and test(20%) sets for training and evaluation respectively.

#### Results

The performance metrics on the updated dataset for both Allo-Allo and Allo-Allo-Ortho are summarized in **Table A1**. The performance results suggest that incorporating orthosteric site information does not enhance—but rather impairs—the prediction of allosteric sites in our framework. One possible explanation is that focusing solely on orthosteric-allosteric interactions might overlook other critical residue-residue relationships that contribute to allosteric mechanisms. By considering all possible residue interactions, the original Allo-Allo model leverages a more comprehensive representation of the protein’s conformational dynamics.

**Table A1:** Performance comparison of Allo-Allo with and without orthosteric site information on proteins with known orthosteric sites.

| Model | AUPRC | AUROC | Acc. | Prec. | Recall | F1 |
| --- | --- | --- | --- | --- | --- | --- |
| Allo-Allo (without orthosteric sites) | **0.79** | **0.96** | **0.98** | **0.85** | **0.64** | **0.73** |
| Allo-Allo-Ortho (with orthosteric sites) | 0.62 | 0.88 | 0.98 | 0.84 | 0.47 | 0.60 |

### C Allo-Allo-AlphaMissense Weighted Risk-Score

Let *M*_*ambiguous*_ and *M*_*pathogenic*_ denote the number of ambiguous and pathogenic mutations respectively for a particular residue. We define the weighted risk-score as

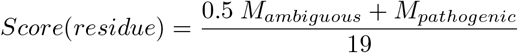

## Notes

### Competing Interest Statement

The authors have declared no competing interest.

